# STAT3 phosphorylation in the rheumatoid arthritis immunological synapse

**DOI:** 10.1101/2025.01.20.633875

**Authors:** Hila Novak Kotzer, Jesusa Capera, Ashwin Jainarayanan, Viveka Mayya, Alexandra Zanin-Zhorov, Salvatore Valvo, Joanne Macdonald, Peter C. Taylor, Michael L Dustin

## Abstract

Targeting the JAK/STAT pathway has emerged as a key therapeutic strategy for managing Rheumatoid Arthritis (RA). JAK inhibitors suppress cytokine-mediated signaling, including the critical IL-6/STAT3 axis, thereby effectively targeting different aspects of the pathological process. However, despite their clinical efficacy, a subset of RA patients remains refractory to JAK inhibition, underscoring the need for alternative approaches. Here, we identify a novel JAK-independent mechanism of STAT3 activation, which is triggered by the formation of the immunological synapse (IS) in naïve CD4+ T cells. Our data demonstrates that Lck mediates the TCR-dependent phosphorylation of STAT3 at the IS, highlighting this pathway as a previously unrecognized hallmark of early T cell activation. Furthermore, we show that the synaptic Lck/TCR-STAT3 pathway is compromised in RA. This discovery highlights a new therapeutic target for RA beyond JAK inhibitors, offering potential avenues for treating patients resistant to current therapies.

## 1. INTRODUCTION

Rheumatoid arthritis (RA) is an autoimmune disease characterized by chronic inflammation of the joints that can lead to irreversible disability. Genetic risk loci include HLA-DR4, cytokine signaling, co-stimulation and inflammatory pathways [1]. The prominence of HLA-DR4 and distinctive peripheral helper T cells implicates the CD4+ T cell immunological synapse (IS) with antigen presenting cells in RA pathology [2, 3]. Environmental factors such as oxidative stress and alterations in DNA methylation also contribute to risk of developing RA [4]. Post-translational modifications, including citrullinated neo-antigens that initiate an adaptive immune response with autoantibody generation, are also a prominent feature, although not always observed [5]. The disease site in the joint is the synovium, which lines the joint capsule and produces the lubricating synovial fluid. Synovial inflammation in RA is characterized by CD4+ T cell and monocyte infiltration, inflammatory fibroblast proliferation and disorganization of the lining layer macrophages. Autoantibodies are assumed to contribute to the pathology based on the success of anti-B cell therapies. Synovitis can persist and eventually cause cartilage and bone destruction, a process perpetuated by the pro-inflammatory cytokine milieu in the joints [6].

Timely and efficient treatment of RA is essential to halt disease progression and to prevent irreversible joint damage. Current treatment for RA includes different classes of disease-modifying anti-rheumatic drugs (DMARDs). Conventional synthetic-DMARDs (cs-DMARDs), like methotrexate, are usually first-line treatments. In the late 90’s, a new family of biological-DMARDs (b-DMARDs) were developed. b-DMARDs are engineered monoclonal antibodies and receptors that target key mediators of RA-associated inflammation. These include recombinant CTLA-4 [7] targeting the RA IS [8], B cell depleting anti-CD20 antibodies [9, 10] to eliminate the major synaptic partner of CD4+ T cells and TNF [11] and IL-6 [12-14] inhibitors targeting key soluble mediators. b-DMARDs are usually administered to patients resistant to methotrexate and shifting from one b-DMARD class to another is common in patients also refractory to some types of b-DMARDs [15].

The latest generation of DMARDs are the Janus kinase (JAK) inhibitors, also known as targeted synthetic-DMARDs (ts-DMARDs). JAKs are cytokine receptor proximal mediators of several cytokine signalling pathways, and therefore mediate different aspects of RA pathophysiology, including autoantibody production in B cells, synovial inflammation, and differentiation of naïve T-cells to helper T-cells. JAK inhibitors have proven their efficacy in RA treatment in patients resistant to methotrexate or to b-DMARDs [16-18].

JAKs are intracellular kinases that associate to specific cytokine receptors and phosphorylate when the ligand-receptor binding occurs. In turn, JAKs activate and phosphorylate the signal transducer and activator of transcription (STAT) transcription factors, which then translocate to the nucleus and regulate gene expression. There are four JAK (JAK1-3 and TYK2) and seven STAT (STAT1-6, including STAT5A and STATB) isoforms. The combination of different expression levels and binding affinities between the cytokine receptor, the JAKs and the STATs involved in a signalling pathway, determines the specific function for each cytokine [19, 20]. Therefore, JAK inhibitors control RA progression at different levels by simultaneously inhibiting several cytokines, such as IL-6 or interferons.

However, despite the efficiency of JAK inhibitors and b-DMARDs, there are still many patients that do not respond to current therapies [6]. Thus, it is essential to further investigate the mechanisms of DMARD unresponsiveness or de-sensitization. Studies on viral transformation show that Src family kinases such as Lck can phosphrorylate and induce DNA binding activity of STAT3 [21], suggesting that STAT3 could be phosphorylated in other contexts where Lck is activated. Here, we identify Lck-dependent STAT3 phosphorylation in CD4^+^ T-cell IS, which bypasses the canonical IL-6-JAK1/2 dependent pathway. In addition, we show that CD4^+^ T-cells from RA patients display more promiscuous STAT3 phosphorylation in the IS, which is less dependent upon Lck. This finding resonates with prior observations that constitutive phosphorylation of STAT3 in CD4^+^ T-cells correlates with RA disease activity [22-24] and that knocking-out STAT3 reduces disease progression in animal models [25]. Therefore, our data describes a novel paradigm for opportunistic STAT3 phosphorylation in the IS as a mechanism through which some RA patients might be refractory to available DMARDs.

## 2. MATERIALS AND METHODS

### 2.1 Isolation of primary human CD4+ T-cells

Primary human CD4+ T-cells were isolated from leukocyte cones provided by UK National Health Service Blood and Transplant, using the negative selection kits RosetteSep™ Human CD4+ T-Cell Enrichment Cocktail (StemCell Technologies). To obtain primary CD45RA+CD4+ T-cells, enriched CD4+ T-cells were further purified using EasySep™ Human “Naïve” CD4+ T-cell Isolation Kit (StemCell Technologies), which enriches naïve and TEMRA CD4+ subsets. Human T-cells were cultured at 37ºC, 5% CO2, in RPMI 1640 medium (Life Technologies) supplemented with 10% FCS, 4 mM L-glutamine, 1% Penicillin-Streptomycin (Gibco), 1% Non-Essential Amino Acids Solution (Thermofisher Scientific), 10mM HEPES (Life Technologies). Informed written consent was obtained prior to obtaining the blood samples as described in the protocol that has obtained approval of the regional ethics committee (REC 07/H0706/81), in accordance with the Declaration of Helsinki.

### 2.2 Preparation of Supported Lipid Bilayer (SLB) and immunocytochemistry

To prepare SLBs [26], clean room grade glass coverslips (Nexterion) were plasma cleaned and mounted onto six-channel chambers (Ibidi). Small unilamellar liposomes were prepared using 1,2-dioleoyl-sn-glycero-3-phosphocholine (Avanti Polar Lipids Inc.) supplemented with 12.5% 1,2-dioleoyl-sn-glycero-3-[(N-(5-amino-1-carboxypentyl) iminodiacetic acid) succinyl]-Ni (Avanti Polar Lipids Inc.). Channels in Ibidi chamber were filled with liposome suspension, blocked with 1% BSA in buffered saline and and washed with 0.1% BSA in buffered saline. SLB were then incubated with the indicated mix of His-tagged proteins in 0.1% BSA in buffered saline to achieve the desired density of molecules on the SLB: UCHT1 recombinant Fab-6His (anti-CD3ε) (30 molecules/ μm^2^) and ICAM1 (200 molecules/μm^2^), without CD80 (200 molecules/μm^2^) as indicated. We determined the amount of the his tagged proteins needed to achieve a given density by quantitative flow cytometry on bead SLBs. His-tagged proteins were produced in 293T cells, purified by Ni^2+^ affinity and size exclusion chromatography and were conjugated with the AlexaFluor dyes of interest (405 or 488). To generate the synapses, cells were exposed to the SLBs at 37 ºC for 20 min, fixed with 4% PFA in PHEM buffer (10 mM EGTA, 2 mM MgCl2, 60 mM Pipes, 25 mM HEPES, pH 7.0) and washed with PHEM buffer to preserve LFA-1-ICAM-1 interactions [26]. For immunofluorescence, cells were permeabilized with 0.1% Triton X-100 in PHEM buffer, blocked in PHEM buffer with 1% BSA and incubated with the primary antibody: anti-pSTAT3 (clone 13A3-1 Biolegend or Cat no 9145 Cell Signaling Tech); anti-Lck (Cat no 2787 Cell Signaling Tech.); anti-pZAP70 (Cat no 2701 Cell Signaling Tech); or anti-PKCθ (Cat no 9376 Cell Signaling Tech) for 1h at room temperature (RT) or overnight at 4^°^C in PHEM buffer with 0.1% BSA. When required, samples were further incubated with a secondary antibody for 1h at RT, washed and then imaged. Note that the secondary antibody could not be anti-mouse due to the UCHT1 Fab containing moue H and L chain sequences.

### 2.3 Total internal reflection fluorescence microscopy (TIRFM), Confocal Microscopy and image analysis

TIRF imaging was performed on an Olympus IX83 inverted microscope with a TIRF module. The instrument was equipped with an Olympus UApON 150x TIRF N.A 1.45 objective, 405 nm, 488 nm, 568 nm and 640 nm laser lines and Photometrics Evolve delta EMCCD camera. Confocal microscopy was carried out on a Zeiss 980 LSM using a 63x oil 1.40 NA objective and appropriate factory-set filters and dichroic mirrors for different fluorophores. Acquisition settings were maintained constant throughout the experiment. Images visualization and analysis was performed using Fiji (ImageJ).

### 2.4 Intracellular Flow Cytometry

To assess levels of intracellular phosphorylated proteins, cells were fixed in 2% PFA for 15 min, washed and gently resuspended in ice-cold 90% methanol and incubated for a minimum of 3 h at -20^°^C. Next, cells were washed and incubated with the corresponding antibody in 0.5% BSA PBS solution for 1h. Samples were washed and analysed using the LSR II Flow Cytometer (BD Biosciences). Data were analysed using FlowJo version 10.7.1.

### 2.5 Western Blotting

Analysis of pSTAT3 vs STAT3 was performed by reprobing the same gels and running parallel blots on duplicated lanes. Cells were lysed in RIPA buffer supplemented with protease and phosphatase inhibitor cocktail (CST Technologies). Equal amounts of lysate were loaded on an SDS-PAGE gels, in some cases in duplicate, and transferred to nitrocellulose membranes. The membranes were blocked, the duplicated lanes when relevant, probed with anti-phospho(Y705)STAT3 (Cat no 9131 or Cat no 4113 Cell Signaling Tech.), anti-Lck (Cat no 2787 Cell Signaling Tech.), anti-phospho PLCg1 (Cat no 14008 Cell Signaling Tech) overnight, washed, and stained with IR dye labeled secondary antibodies from Li-Cor, Inc. (Lincoln, NE). Anti-STAT3 (Cat no 12640S or 9139, Cell Signaling Tech) and anti-beta actin (Cat no 3700S Cell Signaling Tech) were used as loading-controls. Immunoreactive protein bands were visualized using an Odyssey Infrared Imaging system.

### 2.6 Gene disruption by CRISPR/Cas9 on CD45RA+CD4+ T-cells

Gene disruption in primary CD45RA+CD4+ T-cells was performed by transfection with in vitro-prepared Cas9 ribonucleoprotein (RNP) complexes. Briefly, Gene-specific Alt-RCRISPR-Cas9 guide-RNAs (IDT) were incubated with Alt-R tracrRNA (IDT) in nuclease-free duplex buffer (IDT) at 95°C for 5 min and resultant duplex allowed to cool to room temperature. Next, Alt-R S.p Cas9 Nuclease V3 (IDT) and duplexed guide-RNA were mixed in nuclease-free duplex buffer and incubated at 37°C for 15 min. Finally, Alt-RCas9 Electroporation Enhancer (IDT) was added to the RNP solution, and the whole mix then added to 1.5 × 10^6^ CD45RA+CD4+ T-cells. Cells had been previously washed and resuspended with room-temperature OptiMEM. The cell-RNP mix was transferred to a Gene Pulser cuvette (BioRad) and pulsed for 2 ms at 300 V in an ECM 830 Square Wave Electroporation System (BTX). Cells were then immediately transferred to growing media and cultured for 3 days before confirming protein expression and running experiments.

### 2.7 Single-cell RNA sequencing (scRNA-seq)

The raw scRNA-seq data was downloaded from Gene Expression omnibus (GEO) under accession number GSE126030, specifically utilizing four samples corresponding to PBMCs (Peripheral Blood Mononuclear Cells) from this accession code: GSM3589418, GSM3589419, GSM3589420, and GSM3589421. T-cells were isolated using magnetic negative selection for CD3+ T-cells (MojoSort Human CD3+ T-cell Isolation Kit; BioLegend), and dead cell were removed using a kit from Miltenyi Biotec. 0.5 to 1 million CD3+ enriched cells were cultured from each donor tissue for 16 hours at 37°C in complete medium, with the option for TCR stimulation or without. T-cells from these PBMCs were activated using Human CD3/CD28 T-cell Activator (provided by STEMCELL Technologies) and compared to resting T-cells cultured in standard media. The fastq files obtained from GEO were processed using the Cell Ranger (v3.1.0) counts pipeline. The sequencing reads were aligned to Human GRCh38-3.0.0 and the cell count matrices were obtained. These filtered gene–cell matrices were then read into the Seurat v4.0 pipeline implemented in R for downstream analysis. We followed the standard Seurat pre-processing workflow with QC based cell filtration, data normalization and scaling. We further identify major cell types with highly variable genes and performed Jackstraw analysis to isolate significant principal components. Cells of low quality, defined as those with fewer than 200 unique feature counts and cells with a mitochondrial count exceeding 5%, were excluded from the analysis. The UMAP embeddings were then computed using the isolated PCs and known marker genes were used to characterize cell types.

### 2.8 Statistics

All statistical tests were performed using GraphPad Prism 9 software. Student’s t test, Mann-Whitney test and Kruskal-Wallis test were used for statistical analysis and p<0.05 was considered statistically significant. All data is representative from at least 3 independent human blood donors.

## 3. RESULTS

### 3.1 STAT3 is activated at the immunological synapse of total CD4+ and CD45RA+CD4+ T-cells from healthy individuals

STAT3 has pleiotropic roles in CD4+ T-cells, but its activity is typically associated with responses to a number of cytokines including IL-6 and IL-10. To study STAT3 signaling in human CD4+ T-cells in response to TCR signaling in the IS, we used a supported lipid bilayer (SLB) system containing ICAM-1 and anti-CD3, and CD4+ T-cells freshly isolated from peripheral blood (Fig.1A) [27]. In this system, TCR signaling is initiated and sustained in peripheral microclusters and terminated in the IS center [28]. We found that phosphorylated STAT3 (pSTAT3) was polarized at the IS (Fig.1B-C). TCR engagement in total CD4+ T-cells increases phosphorylation of STAT3, based on a 29% increase in fluorescence intensity in microclusters on anti-CD3 + ICAM-1 SLBs, when compared to ICAM-1 only SLBs (Fig.1C). Since pSTAT3 detection in CD4+ T-cells containing naïve, memory and regulatory T-cell was variable, we purified CD45RA+CD4+ T-cells, which were enriched for naïve T-cells, and found that pSTAT3 at the IS was lower, but more uniform (Fig. 1D, Fig.1-Suppl.1). We further demonstrated that treatment with anti-CD3 or anti-CD3/CD28 activation beads up-regulates the whole-cell pSTAT3, in both CD45RA+CD4+ and CD4+ T-cells (Fig.1E-G). CD28 co-stimulation plays an important role in T-cell priming [29, 30]. Thus, we compared the ability of CD45RA+CD4+ T-cells to activate STAT3 at the IS in the presence of CD80 in the SLB. We did not detect a significant difference in pSTAT3 at the IS in the presence or absence of CD80 in the SLB. As a positive control for CD80 activity, we demonstrated that CD80 in the SLB increased PKC-θ recruitment to the IS, validating the activity of CD80 on the SLB (Fig.1-Suppl.2A-B). Finally, we tested the activation of whole-cell STAT3 at different time points, from 20 min and up to 2 hours and found that the signal accumulated with time (Fig. 1G, Fig.1-Suppl2C).

**Figure 1.**
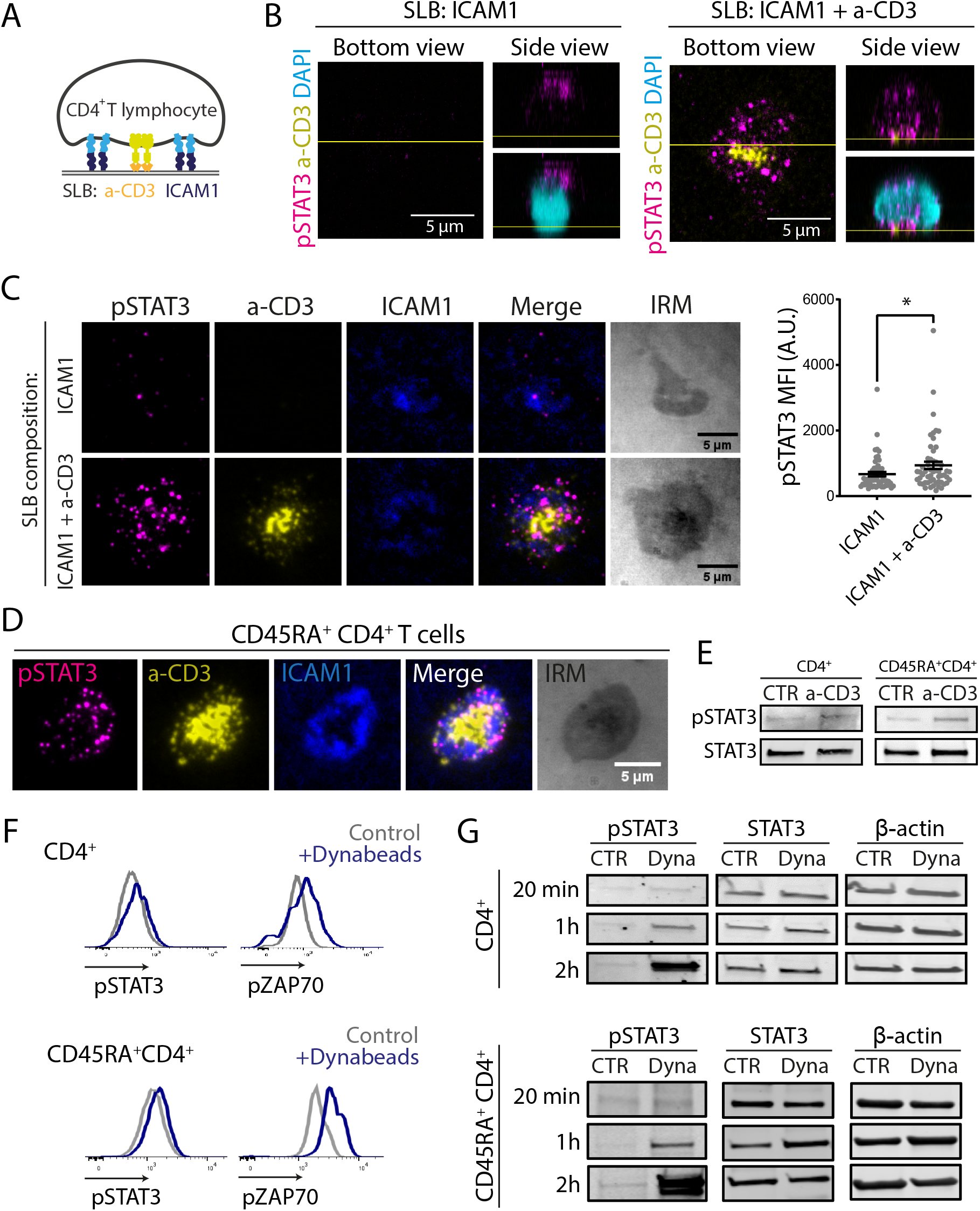
STAT3 is activated in CD4+ and CD45RA+CD4+ T-cells after TCR stimulation. **(A)** CD4+ T-cells are exposed to the SLB, containing His-tagged anti-CD3 and ICAM-1. **(B)** Representative confocal 3D stacks showing CD4+ T-cells forming a contact with SLBs. Samples stained for pSTAT3. **(C)** Representative TIRF images of CD4+ T-cells forming a contact with SLBs. Plot shows pSTAT3 mean fluorescence intensity (MFI). Data is the mean +/-SE (n=55 cells from 3 independent donors); *p<0.05, Mann-Whitney test. **(D)** Representative TIRF images of CD45RA+CD4+ T-cells introduced into SLBs containing anti-CD3 and ICAM-1 and stained for pSTAT3. **(E)** CD4+ and CD45RA+CD4+ T-cells were activated (a-CD3) or not (CTR) with anti-CD3 coated beads for 20 min and total STAT3 and pSTAT3 protein expression were measured by Western blot. **(F)** CD4+ and CD45RA+CD4+ T-cells were activated (Dyna) or not (CTR) with Dynabeads. After 20 min, pSTAT3 expression was measured by Flow Cytometr. pZAP70 was used as a positive control for TCR activation. **(G)** CD4+ and CD45RA+CD4+ T-cells were activated (Dyna) or not (CTR) with Dynabeads for the indicated timepoints before protein extraction and analysis by Western blot. STAT3 and β-actin were used as loading controls for Western Blots on samples run in parallel.

### 3.2 STAT3 activation at the immunological synapse is Lck dependent

We next examined the effect of Lck inhibitor on STAT3 phosphorylation at the IS of CD45RA+CD4+ T-cells. We used the selective Lck inhibitor (A-770041, Axon Medchem), and pre-treated CD45RA+CD4+ T-cells before introducing the cells to SLBs containing anti-CD3 and ICAM-1. We found that the Lck inhibitor abrogated the phosphorylation of STAT3 at the IS (Fig. 2A) and confirmed reduction of pSTAT3 through Lck inhibition through immunoblotting on the whole cell lysates (Fig. 2B, Fig. 2-Suppl. 2). To further confirm Lck-dependent STAT3 phosphorylation, we knocked-out LCK expression in CD45RA+CD4+ T-cells using the CRISPR/Cas9 technology. Phosphorylation of STAT3 at the IS was significantly reduced in LCK KO cells when compared to the CD19 KO control (Fig. 2C). LCK KO cells also showed reduced whole-cell pSTAT3 after Dynabead stimulation (Fig. 2D).

**Figure 2.**
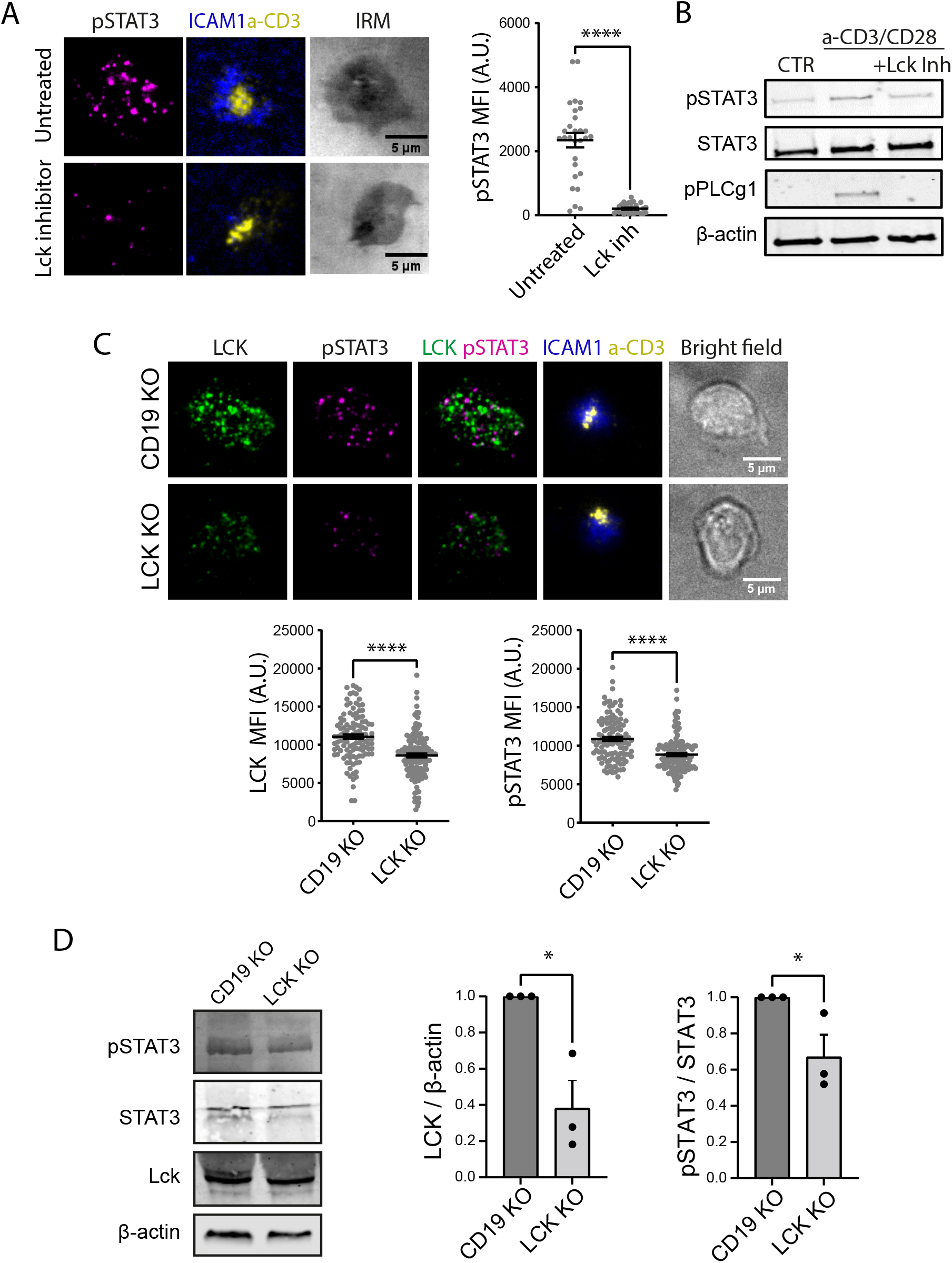
TCR-dependent activation of STAT3 is mediated by Lck. **(A)** (Left) TIRF images of CD45RA+CD4+ T-cells exposed to SLBs (ICAM1 and a-CD3). Cells were pre-treated, or not, for 30 min with Lck inhibitor (A-770041, Axon Medchem, 10µM). (Right) Quantification of pSTAT3 MFI. Data is the mean +/-SE (n=30 cells); ****p<0.0001, Mann-Whitney test. **(B)** Western Blot analysis of CD45RA+CD4+ T-cells treated or not (CTR) with beads coated with anti-CD3/CD28 for 20 min. When indicated, cells were treated with Lck inhibitors. pSTAT3 and total STAT3 immunoblots were run in parallel and analyzed in relation to loading controls **(C)** (Top) TIRF images of LCK Knock-out (KO) CD45RA+CD4+ T-cells forming synapses with SLBs (ICAM1 and a-CD3). CD19 KO, negative control. Samples were stained for Lck and pSTAT3. (Bottom) Quantification of Lck and pSTAT3 MFI. Data is the mean ± SE (n>90 cells for 3 donors); ****p<0.0001, Mann-Whitney test. **(D)** KO CD45RA+CD4+ T-cells were treated with Dynabeads for 20 min and total protein expression was analyzed by Western Blot. Plots show quantification of total Lck protein expression normalized by β-actin and total pSTAT3 protein expression normalized by total STAT3. Data was relativized to CD19 KO. Data are the mean ± SE from 3 donors; *p<0.05, one-tailed Student’s t-test.

### 3.3 Jak1 inhibitors do not inhibit STAT3 activation at the IS of CD45RA+CD4+ T-cells

To further confirm the critical role of Lck in STAT3 phosphorylation at the IS of CD45RA+CD4+ T-cells, we used two different Jak1 inhibitors. Ruxolitinib, a Jak1/Jak2 inhibitor, and filgotinib, a selective Jak1 inhibitor [31] (Fig. 3A). We first validated the activity of both inhibitors by treating purified CD45RA+CD4+ T-cells with just IL-6, or with IL-6 in the presence of each inhibitor (Fig. 3B, Fig. 3-Suppl1). This confirmed that IL-6-dependent STAT3 activation was diminished in the presence of either ruxolitinib or filgotinib, both at the whole cell level (Fig. 3B) and at the IS (Fig. 3-Suppl 1). Next, we evaluated the effect of both inhibitors on STAT3 activation at the IS of CD45RA+CD4+ T-cells. Cells were pre-treated with either of the Jak inhibitors or the carrier for 45 min. Then, cells were exposed to SLBs containing anti-CD3 and ICAM-1 to form IS, fixed and analyzed for pSTAT, or the inhibitors were washed-out and replaced by fresh media right before introducing the cells into the SLBs to form IS. Ruxolitinib did not generate a significant effect compared to the carrier pre-treatment, but reduced the amount of pSTAT3 by 19 % at the IS when compared to the washout condition (Fig 3C). No difference in pSTAT3 levels were observed for filgotinib treatment under any condition (Fig 3D). This data suggests that Jak1/2 have a small, marginally significant effect on STAT3 phosphorylation at the IS compared to the greater and more consistent effect of Lck inhibition.

**Figure 3.**
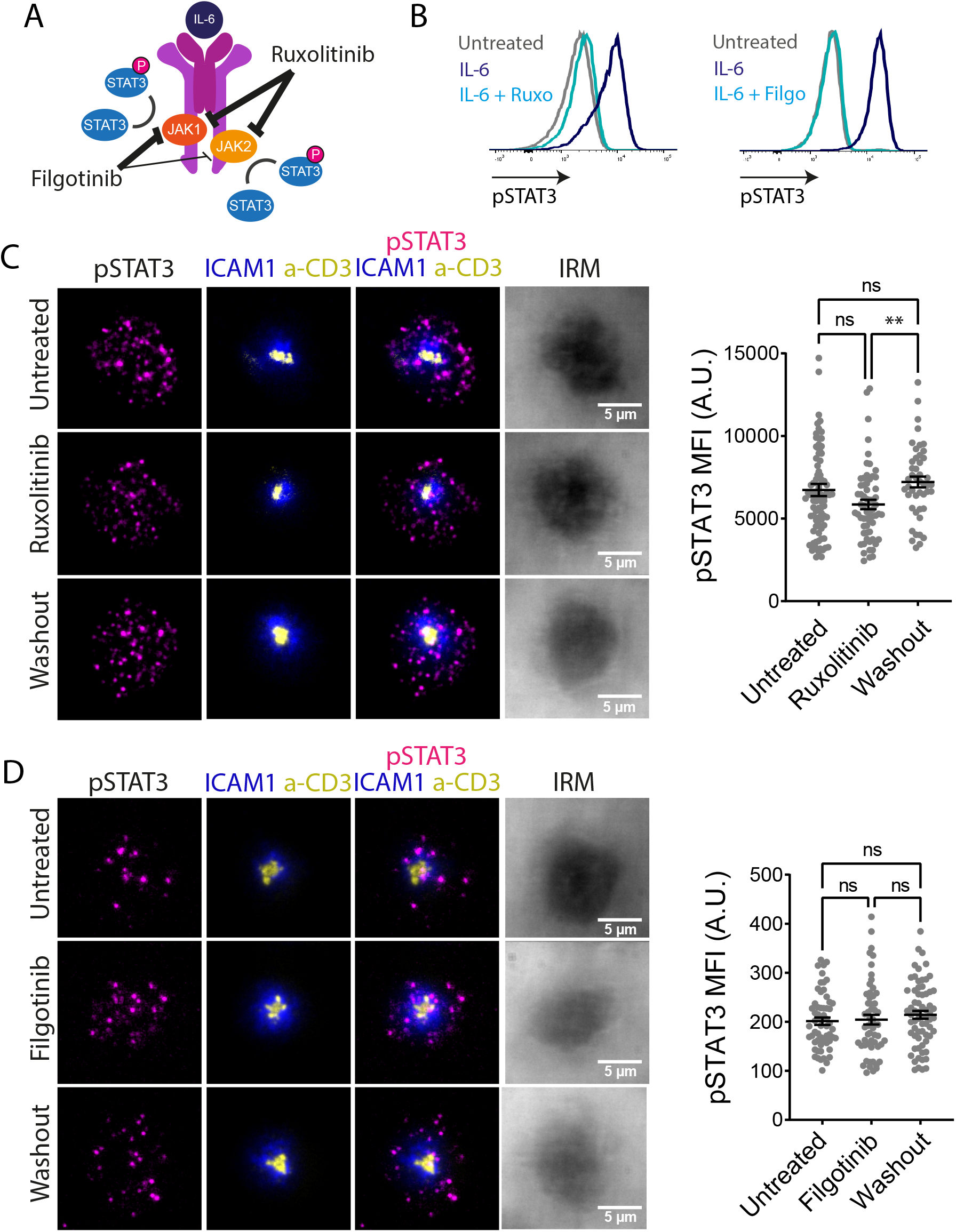
TCR-dependent activation of STAT3 is not mediated by Jak. **(A)** Cartoon representing the IL-6 receptor signaling pathway. Ruxolitinib inhibits both Jak1 and Jak2, filgotinib preferentially inhibits Jak1. **(B)** CD4+ T-cells were treated or not with 25 ng/ml of IL-6 for 15 min and total pSTAT3 levels were assessed by Flow Cytometry. When indicated, cells were pre-treated for 45 min with 10 µM of the indicated Jak inhibitor (Left, ruxolitinib. Right, filgotinib). **(C-D)** (Left) Representative TIRF images of CD45RA+CD4+ T-cells forming synapses with SLBs (ICAM1 and a-CD3). Cells were pre-treated or not (untreated) for 45 min with 10 µM of ruxolitinib (Top) or filgotinib (Bottom). When indicated, the inhibitor was washed-out before SLB exposure. Cells were fixed and permeabilized after 20 min of SLB exposure and pSTAT3 expression was analyzed. (Right) Plots show quantification of pSTAT3 MFI. Data is the mean ± SE (n>50); ns, not significant; **p<0.01, Kruskal-Wallis test.

### 3.4 TCR engagement upregulates STAT3 related genes

We analysed publicly available scRNAseq datasets in order to determine the expression levels of STAT3 in different subsets of CD4+ T-cells from peripheral blood of healthy individuals in a resting state or activated using human CD3/CD28 T-cell activator (provided by STEMCELL Technologies). We identified naïve, effector, memory, senescent, exhausted and FOXP3+ subsets based on unbiased clustering and signature genes [32] (Fig. 4A, Fig. 4-Suppl.1). As expected, we identified a reduced naïve T-cells proportion following activation (Fig. 4A). In addition, a feature plot (Fig. 4A) of the STAT3 expression among activated and resting cells revealed that STAT3 was enriched in the activated T-cell population. This conclusion was reinforced by the violin plots (Fig. 4B) of STAT3 expression among all cell types in activated and resting states. This implied that STAT3 enrichment was a universal aspect of activation across all CD4+ T-cell subtypes. Further, the enrichment of TCR related genes in CD4+ T-cell subtypes upon activation was also confirmed; specifically, among the memory, FOXP3 and effector clusters (Fig. 4C). As a negative control, we confirmed that IL-6, IL10, IL11 and related genes that would be likely to activate STAT3 were not upregulated upon TCR stimulation (Fig. 4C and Fig.4-Suppl.2). On the other hand, expression of STAT3 related genes was enhanced upon activation including ZFP36, JUNB, BATF, IL2RA, ICOS, PIM2, SOCS1, SOCS3, IFNG, HIF1A, IRF4 and TNF (Fig. 4C). In contrast to TCR-related and STAT3-related genes, we did not observe much difference between activated and resting cells with respect to IL-6 related genes except for JUNB, SOCS1 and GBP2 as these genes known to be co-regulated by STAT3. IL-6 related genes like CISH, MCL1, PIM1, IFIT1 also showed no difference between activated and resting CD4+ T-cell subtypes. They were indeed less expressed in the activated sample among effector cells. Thus, transcriptional programmes in human CD4+ T-cells activated via the TCR were consistent with TCR dependent STAT3 phosphorylation.

**Figure 4.**
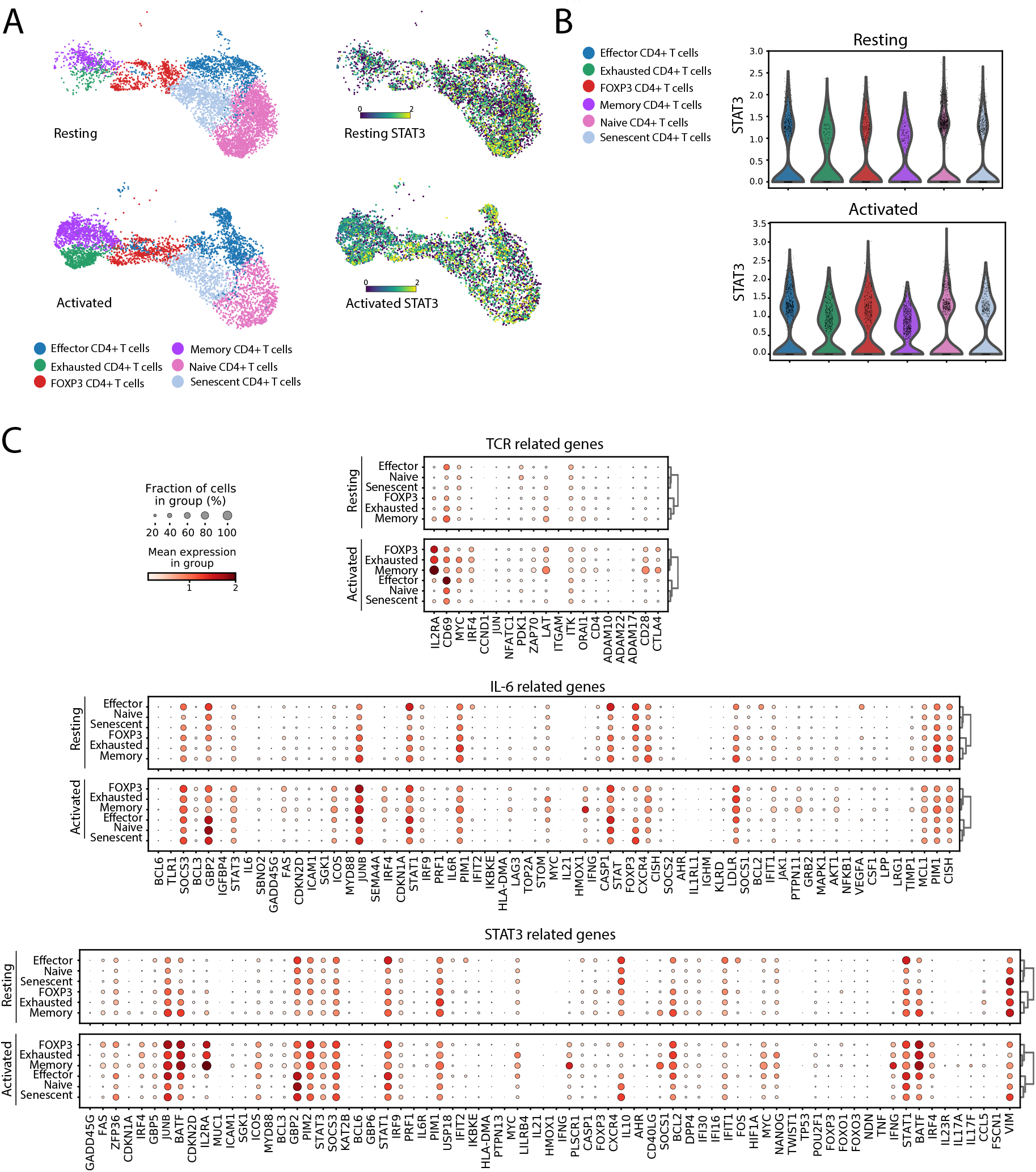
STAT3-related genes upregulate expression in response to TCR engagement. **(A)** (Left) UMAP dimensionality reduction embedding of the activated and resting CD4+ T-cells coloured by orthogonally generated clusters labelled by manual cell type annotation. (Right) A UMAP embedding of scRNAseq dataset of activated and resting CD4+ T-cell population coloured by expression profiles of STAT3. **(B)** Violin plots of the STAT3 expressions across the different cell types in the activated and resting CD4+ T-cell population. **(C)** Dot plots depicting mean expression (visualized by colour) and fraction of cells (visualized by the size of the dot) expressing key genes involved in TCR signaling (Top), IL6 signaling (Middle) and STAT3 regulation (Bottom) in resting and activated CD4+ T-cell subtypes.

### 3.5 RA patient’s CD4^+^ T cells are less dependent on Lck for STAT3 activation at the IS

We next asked if TCR-dependent STAT3 phosphorylation is altered in RA, as dysregulated STAT3 signaling in CD4+ T-cells has been proposed as an early pathophysiological event in RA [33]. We determined the level of STAT3 phosphorylation at the IS without and with Lck inhibition in the IS of CD4+ T-cells from healthy individuals and RA patients (Suppl. Table1). We purified CD4+ T-cells from the peripheral blood of 21 RA patients and 15 healthy controls. We pre-treated the cells with the selective Lck inhibitor, or treated with the carrier, and introduced the cells to SLBs containing anti-CD3 and ICAM-1. We analyzed STAT3 phosphorylation at the IS using TIRFM. We found that the Lck inhibition of pSTAT3 at the IS of CD4+ T-cells from RA patients was impaired compared to healthy control cells (Fig.5, Suppl. Table1). We conclude that TCR-dependent STAT3 phosphorylation in RA patients is significantly less dependent on Lck compared to healthy controls, suggesting a pathological decoupling of STAT3 activation from TCR signaling in RA.

## 4. DISCUSSION

We have provided evidence for TCR driven STAT3 activation at the IS in human CD4+ T-cells (Fig. 1). By using the SLB system, we show polarization of pSTAT3 at the immunological synapse. This constitutes the first observation of STAT3 activation within the synapse. The activation of STAT3 accumulates over time. Through pharmacological inhibition and CRISPR we found that this TCR-driven activation is Lck dependent (Fig. 2). Pharmacological inhibition of Jak1/2 only slightly impaired IS pSTAT3 (Fig. 3). We found that T-cell activation results in increased STAT3 gene expression signatures across CD4+ T-cell subsets. These results are consistent with prior biochemical analysis that Lck can phosphorylate STAT3 [34] and that TCR triggering activates STAT3 in the absence of cytokines [35]. Similarly, we have recently described the TCR-dependent activation of STAT5 at the immunological synapse [36]. The manner in which STAT3 is recruited for phosphorylation in the IS was not determined here. One possible mechanism could involve CXCR4, which can interact with the TCR [37] and recruits STAT3 [38], but a recent study suggested that TCR-CXCR4 interactions would only be transient [39]. A recent study provides compelling evidence that TCR- and Lck-dependent STAT3 activation promotes Th17 differentiation [40]. A key finding in our study relates to a reduced requirement for Lck in TCR dependent STAT3 phosphorylation in the IS of T-cells from RA patients (Fig. 5).

**Figure 5.**
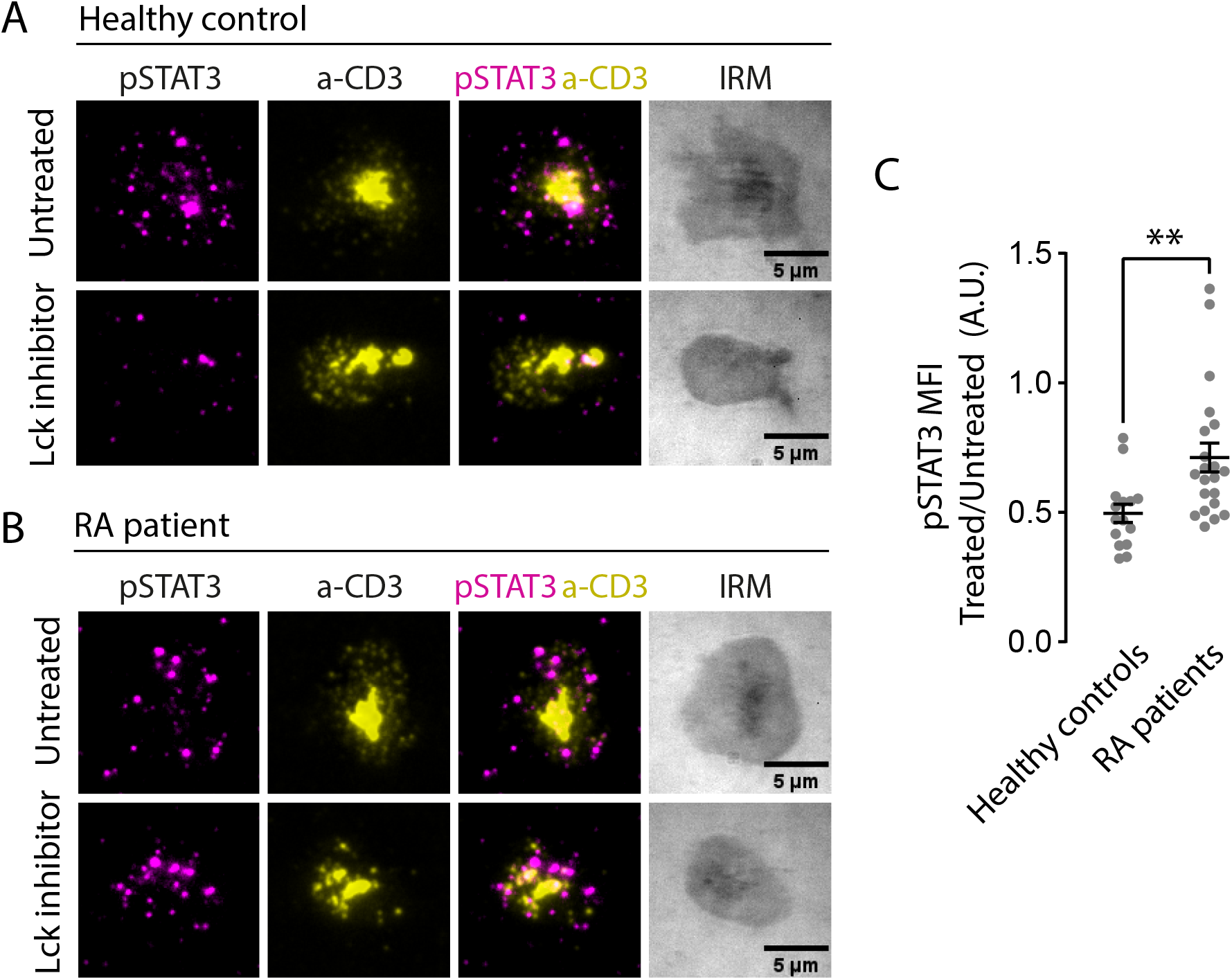
STAT3 activation is less sensitive to Lck inhibition in Rheumatoid Arthritis (RA) patients. **(A-B)** Representative TIRF images of CD4+ T-cells from healthy donors (A) or from RA patients (B) exposed to SLBs (ICAM1 and a-CD3) for 20 min. Samples were then fixed, permeabilized and stained for pSTAT3. When indicated, cells were pre-treated for 30 min with Lck inhibitor (10µM). **(C)** Quantification of pSTAT3 mean fluorescence intensity (MFI) for cells in (A-B). Each dot represents the average MFI of one donor relative to the average MFI of the same donor untreated. Data is the mean +/-SE (n=21 RA patients and n=15 healthy donors); **p<0.01, Mann-Whitney test.

The role of STAT3 in RA has been extensively studied in recent years. Aberrant STAT3 signaling has a well-documented role in sustaining autoimmunity [24, 41-43]. STAT3 has been shown to promote the survival, proliferation and invasion of RA synoviocytes and the production of pro-inflammatory mediators and matrix-degrading enzymes [6]. Moreover, STAT3 activation correlates with the levels of IL-6, a key cytokine in RA, in circulating CD4+ T-cells from RA patients [22, 44]. IL-6 induces hyperactivation of STAT3 in these cells, which may impair the regulation of immune responses. Using Nur77-eGFP reporter in the SKG mice model revealed that the augmented autoreactivity observed was associated with heightened IL-6 receptor signaling, likely contributing to their arthritogenicity [45]. This suggests that STAT3 signaling is dysregulated in RA, and related disease models, even in the absence of external stimuli, and may contribute to the chronicity and severity of the disease.

Our analysis of public scRNA seq on resting and activated human T-cells revealed a significant STAT3 signature, but a limited activation of IL-6 associated genes, which would have been the major candidate to activate STAT3 prior to our findings here (Fig. 4). Pratt et al [46] established a 12-gene ’RA expression signature’ for anti-citrullinated protein autoantibodies (ACPA)-negative RA patient group. This signature was interpreted to implicate an IL-6-mediated STAT3 transcription program in peripheral blood CD4 T-cells of early RA patients. Subsequently, Anderson et al [33] conducted an independent cohort study involving 161 patients, reaffirming the earlier findings. Their research indicated that 11 out of the 12 proposed signature genes exhibited significant differences between RA patients and controls. Of these genes, PIM1, BCL3, and SOCS3, which are regulated by STAT3, showed the most prominent differential regulation. In a meta-analysis of 279 patients, these three genes were found to be primarily responsible for the signature’s ability to discriminate RA patients from healthy controls, irrespective of age, joint involvement, or acute phase response. Furthermore, in vitro studies by Ridgley et al [47] demonstrated that pre-treating CD4+ T-cells with IL-6 led to dysregulated gene expression predisposing to heightened proliferative capacity and the inclination towards Th1-skewing upon subsequent antigen exposure. Taken together, these studies collectively highlight the critical role of STAT3-mediated signaling in the early phases of RA. In this context, our results suggest a link between TCR mediated and cytokine mediated STAT3 signaling that may be altered in RA.

We find that although STAT3 activation at the IS in healthy controls was dramatically reduced by Lck inhibition, in T-cells from active RA patients, Lck inhibitors were less effective in reducing pSTAT3 in the IS (Fig. 5). Because STAT3 activity was shown to be increased in RA patients [24, 44], it is possible that the decreased Lck dependence we observed in the IS reflects opportunistic activation of STAT3 by other kinases in the context of TCR mediated close contacts, even when TCR signaling is disabled. Even when Lck is inhibited, we expect that the formation of actin-dependent TCR microclusters will exclude CD45 [28, 48] and create an environment where other kinases, including Jaks, would be partly protected from dephosphorylation [49]. Further studies are required to test this hypothesis. Cellular and biochemical analyses conducted on human CD4+ T-cells have unveiled irregular TCR signaling among RA patients [50]. CD4+ T-cells isolated from RA patients are hyporesponsive to TCR engagement when tested ex vivo, as demonstrated by their reduced calcium mobilization and interleukin-2 (IL-2) production [51, 52]. This might be attributed, in part, to the decreased expression of TCRζ and linker of activated T-cells (LAT), and changes associated with immune senescence [53-56]. While CD4+ T-cells from individuals with RA exhibit reduced signaling capacity upon in vitro stimulation, they demonstrate hyperproliferation during clonal expansion. Subsequently, these cells differentiate into effector cells that contribute to the progression of the disease. Partial loss of function in the TCR associated kinase ZAP-70 causes autoimmunity in the SKG mouse model for RA. Thus, our results have identified a new alteration in TCR signaling in RA-reduced Lck dependence of STAT3 phosphorylation.

Current targeted therapies for RA block tumour necrosis factor, T-cell costimulation, inflammatory cytokines and B cells [57]. JAK inhibitors are small-molecule oral treatments targeting cytokine receptor signaling [16-19]. Surprisingly, we found that treatment of CD4+ T-cells with selective and orally available Jak1 and Jak2 inhibitors only slightly reduced pSTAT3 in the IS (Fig 3). However, inhibition of Lck, a Src family kinase, resulted in significant inhibition of pSTAT3 in the IS, although this effect was less pronounced in RA patients. Further study is required to determine if the Lck independent component of pSTAT3 activation in RA patient T-cells is Jak dependent or relies on other kinases like Fyn or Abl [58]. The heterogeneity in RA patient responses suggests the potential to use Lck dependence of pSTAT3 in the IS to stratify RA patients [59, 60].

In conclusion, our data demonstrate that STAT3 is activated in the IS. This finding reinforces earlier findings that the TCR can activate STAT3. We show that IS associated pSTAT3 is significantly dependent on Lck and not Jak1/2. The dependence on Lck is reduced in RA, suggesting opportunistic STAT3 activation by other tyrosine kinases with potential to circumvent targeted therapies. These findings suggest a previously uncharacterized paradigm that may be of importance for evaluating new therapeutic approaches for STAT3-related autoimmune diseases and malignancies.

## 5. DATA AVAILABILITY

The data are available from the corresponding author upon reasonable request.

## 6. ACKNOWLEDGEMENTS

We thank E. Kurz, L. Chen, J. Afrose, H. Rada and L. Cifuentes for essential support. We would also like to thank all the anonymized blood donors who contributed to our study. This work was supported by a Cancer Research Institute Irvington fellowship to JC (CRI4503), Cue Biopharma, the Kennedy Trust for Rheumatology Research 202117, the NIH R37 AI043542, the Oxford-BMS fellowship programme and the Wellcome Trust 100262Z/12/Z and 224343/Z/21/Z.

## 8. AUTHOR CONTRIBUTIONS

Conceptualization: HNK, JC, VM, AZZ, MLD; Investigation: HNK, JC, AKJ, VM, AZZ, SV; Formal analysis: HNK, JC, AKJ. Resources: JM, PCT; Writing-original draft: JC, HNK, AKJ, MLD; Writing-review and editing: JC, HNK, AKJ, JM, PCT, MLD

