## Supplementary Material for "STAT3 phosphorylation in the rheumatoid arthritis immunological synapse"

*Figure 1-Suppl 1.* Quantification of pSTAT3 mean fluorescence intensity (MFI) at the synapse of the indicated cells exposed to SLBs (ICAM1 and a-CD3) and imaged by TIRFM.

*Figure 1-Suppl 2. Effect of CD80/CD28 engagement on TCR-dependent STAT3 activation.* **(A)** (Left) Representative TIRF images of CD45RA+CD4+ T-cells exposed to SLBs with the indicated protein composition. Cells were fixed after 20 min of SLB exposure, then permeabilized and stained for PKCΘ. (Right) Quantification of PKCΘ mean fluorescence intensity (MFI). Data is the mean +/- SE (n>25 cells). **(B)** (Left) Representative TIRF images of CD45RA+CD4+T-cells exposed to SLBs with the indicated protein composition. Cells were fixed after 20 min of SLB exposure, then permeabilized and stained for pSTAT3. (Right) Quantification of pSTAT3 mean fluorescence intensity (MFI). Data is the mean +/- SE (n>80 cells). **(C)** CD4+ T-cells and CD45RA+CD4+ T-cells were treated or not (CTR) with beads coated with anti-CD3, anti-CD3/CD28 or Dynabeads as indicated for 2h. pSTAT3 protein expression was then assessed by Western Blotting followed by reprobing with total STAT3 as a loading control.

*Figure 2-Suppl 1. TCR-dependent activation of STAT3 is mediated by Lck.* **(A-B)** Representative TIRF images of CD4+ (A) and CD45RA+CD4+ (B) T-cells exposed to SLBs (ICAM1 and a-CD3). Cells were pre-treated, or not, for 30 min with Lck inhibitor (A-770041, Axon Medchem, 10µM). Samples were fixed and permeabilized after 20 min of SLB exposure and stained for pZAP70. **(C-D)** CD4+ (C) and CD45RA+CD4+ (D) T-cells were treated or not (CTR) with beads coated with anti-CD3 or anti-CD3/CD28 for the indicated timepoints. When indicated, cells were also treated with Lck inhibitors (+Lck inh). pSTAT3 protein expression was analysed by Western Blots on samples run in parallel. STAT3 and β-actin were used as loading controls and pPLCγ1 was used as a positive control of Lck activation.

*Figure 3-Suppl 1. Jak inhibitors effectively inhibit IL-6-induced pSTAT3 at the synapse.* **(A)** Representative TIRF images of CD45RA+CD4+ T-cells forming synapses with SLBs (ICAM1 and a-CD3) after treatment (+ IL-6) or not (control) with the indicated interleukin. **(B)** Cells were treated or not with 25 ng/ml of IL-6 and, when indicated, with 10 µM of the indicated Jak inhibitor (ruxolitinib or filgotinib). Cells were fixed and permeabilized after 20 min of SLB exposure and pSTAT3 expression was analyzed. Plots show quantification of pSTAT3 MFI. Data is the mean ± SE (n>30); ns, not significant; ***p<0.001, ****p<0.0001, Kruskal-Wallis test.

*Figure 4-Suppl 1. Cell type annotations.* **(A)** Heatmaps visualization of differentially expressed genes between different CD4+ T cell subtypes. **(B)** Trackplot illustrating qualitative gene expression signatures of CD4+ T cell subtypes.

*Figure 4-Suppl 2.* Other genes that activate STAT3. Dot plots depicting mean expression (visualized by colour) and fraction of cells (visualized by the size of the dot) expressing key genes that would likely activate STAT3.

*Suppl Table 1.* The clinical and demographic details of the studied RA patients.

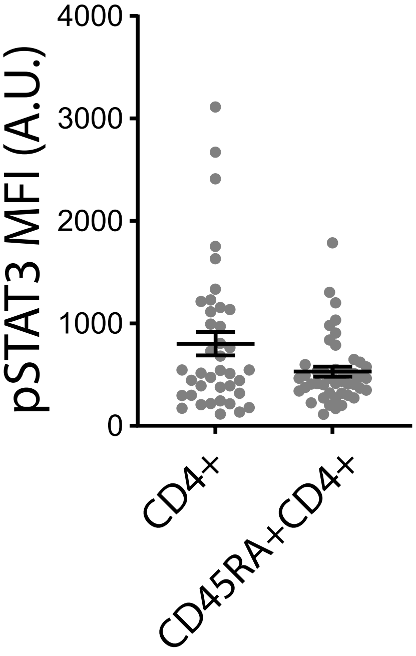


Figure 1 Suppl 1

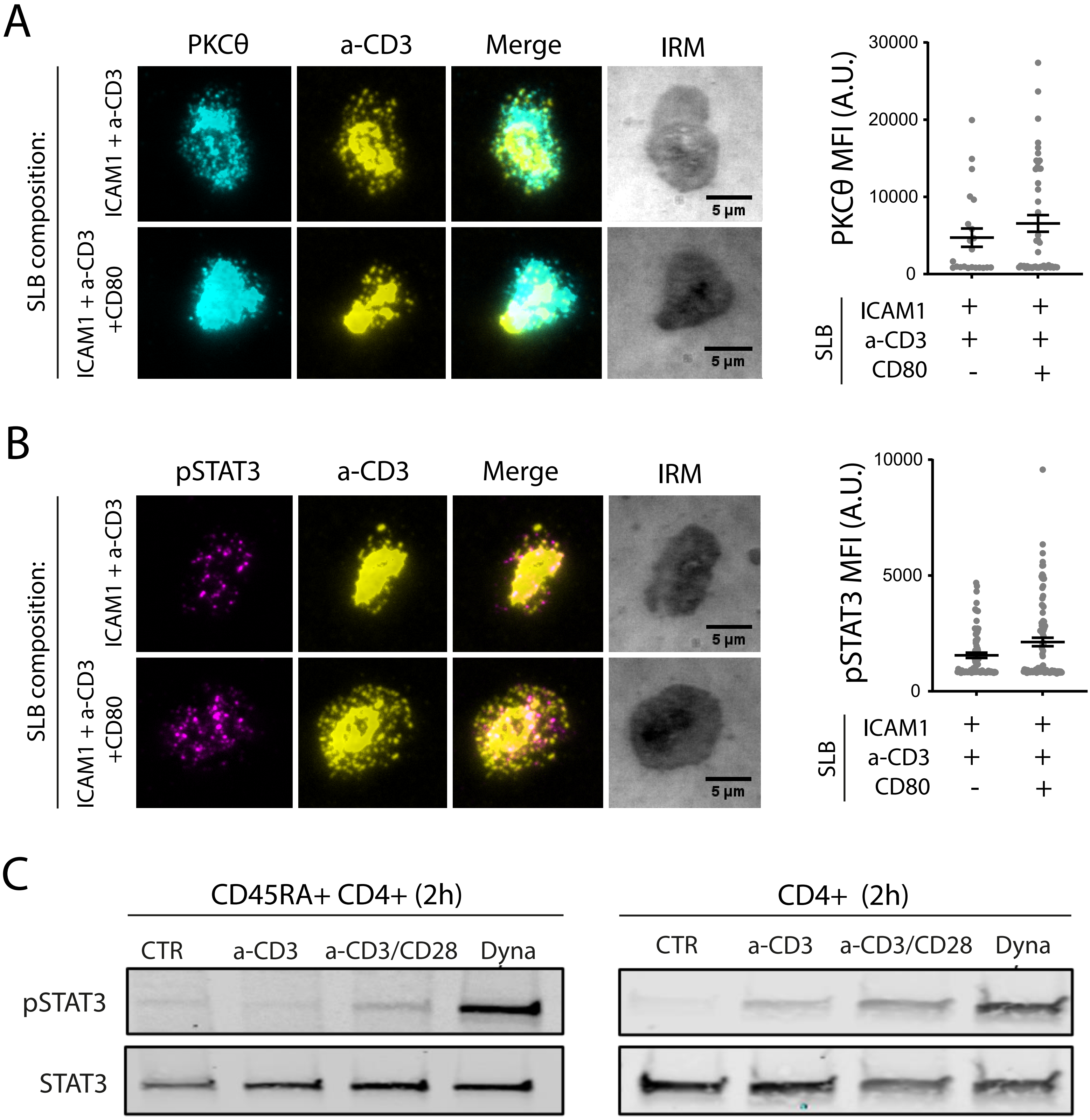


Figure 1 Suppl 2

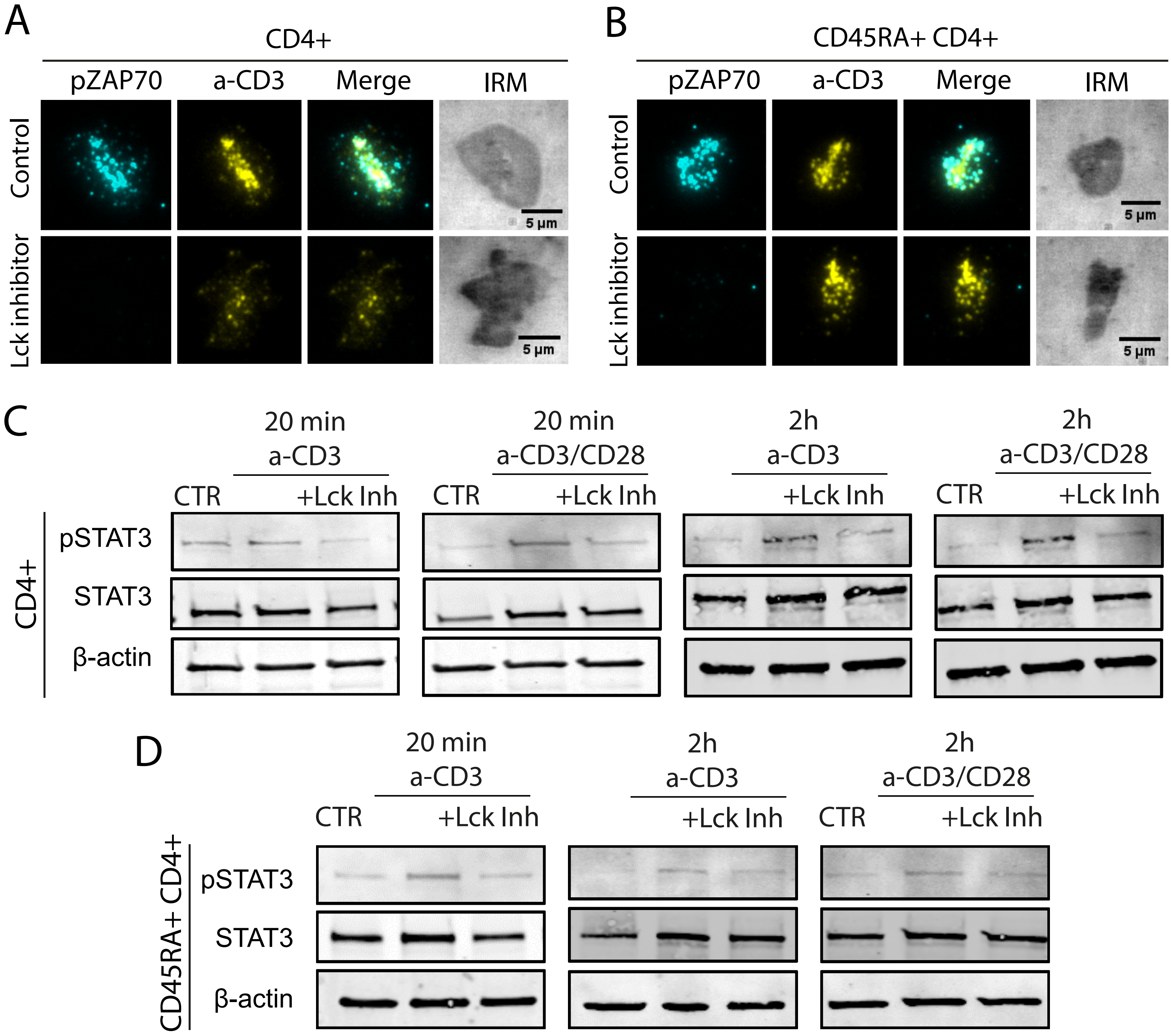


Figure 2 Suppl 1

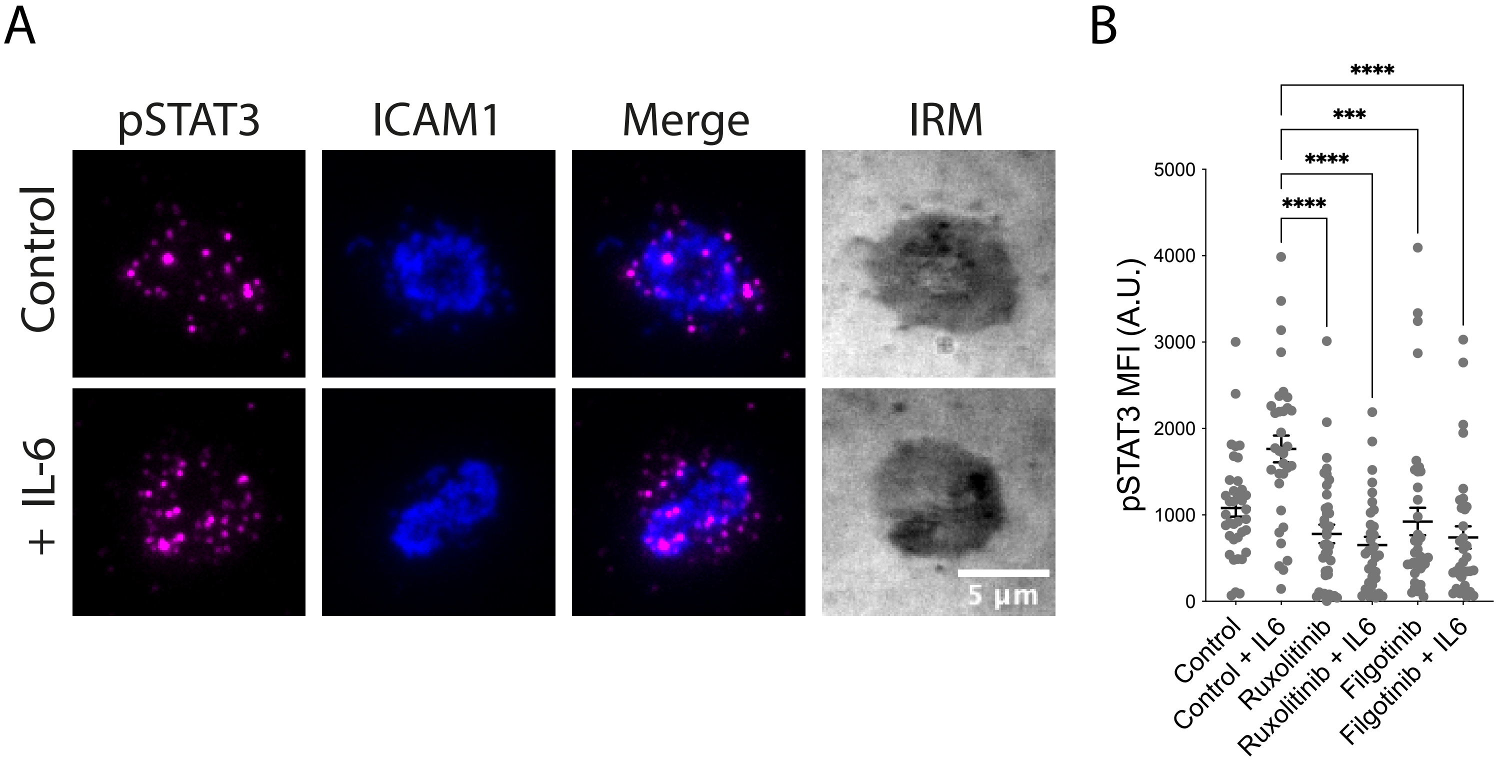


Figure 3- Suppl 1

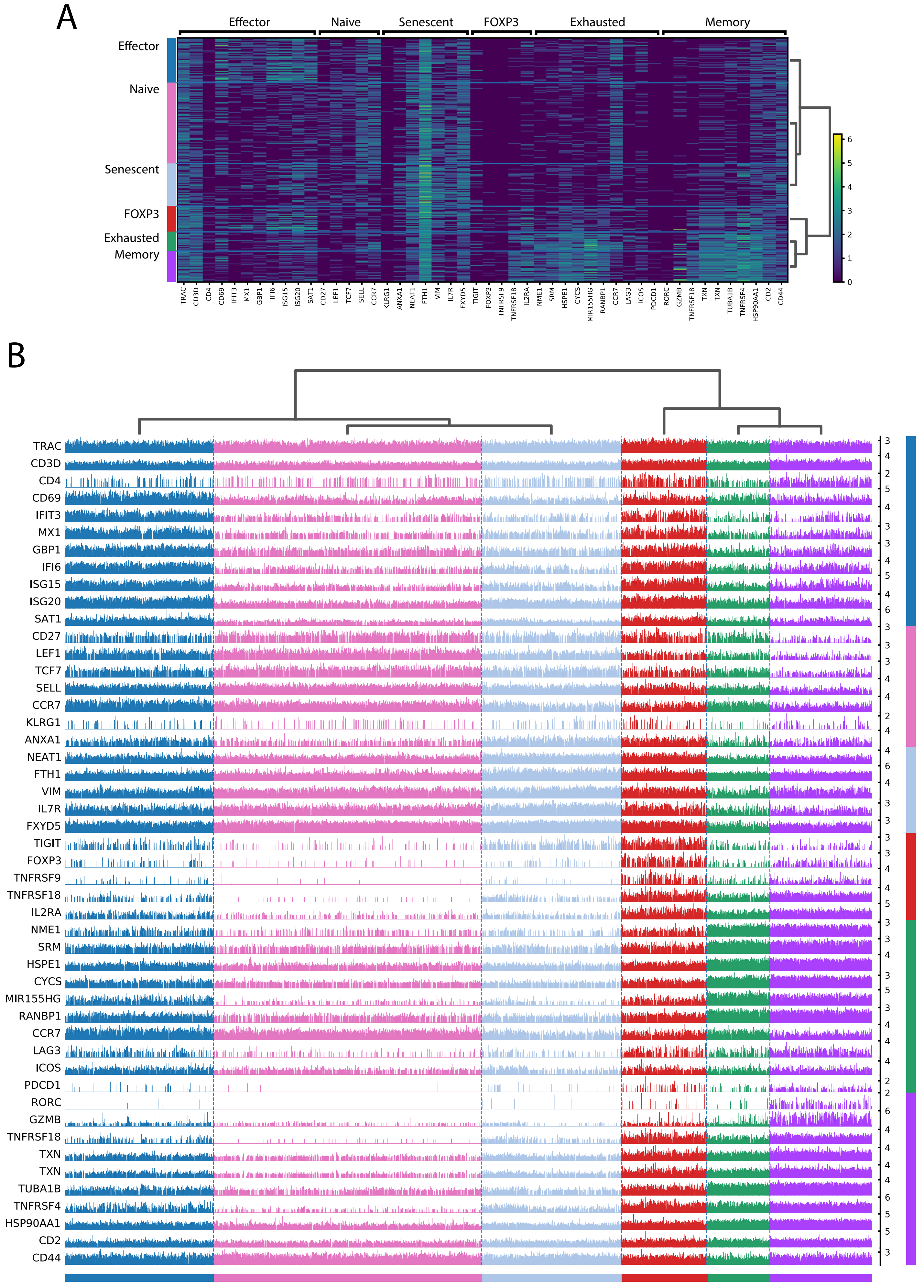


Figure 4 Suppl 1

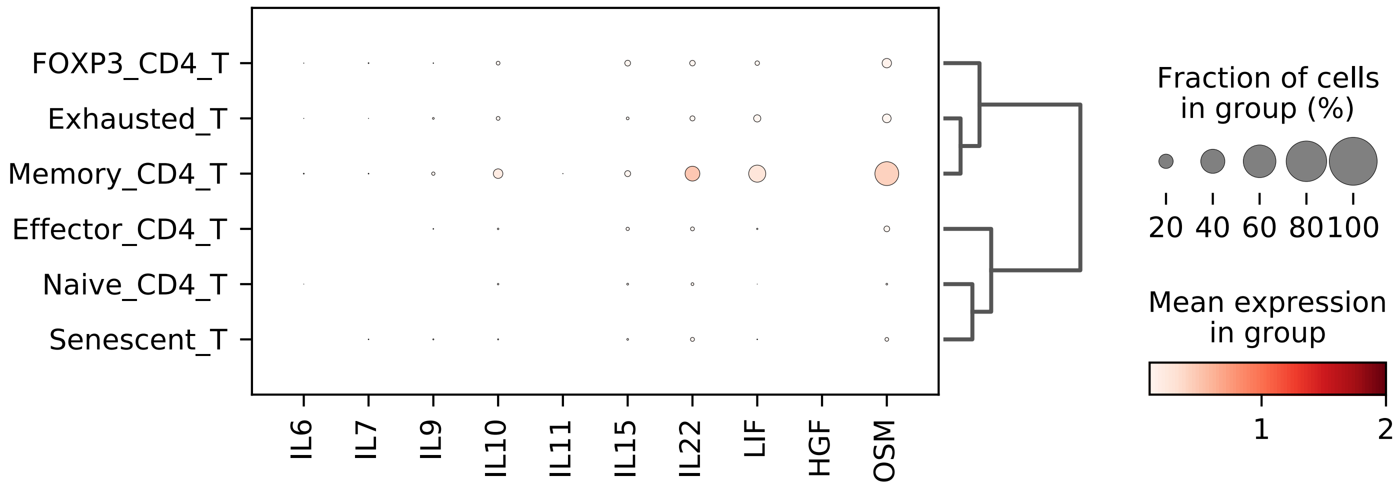


Figure 4 Suppl 2

| study number | cross check of diagnosis | Gender | Age | Current Medication | DAS28-CRP |
|---|---|---|---|---|---|
| 1001 | Rheumatoid arthritis (seronegative, anti-CCP negative) | Female | 55 | Adalilmumab,Methotrexate, Aciclovir | 1.81 low disease activity |
| 1002 | Erosive inflammatory arthritis likely psoriatic with prominent nail ridging and onycholysis throughout. | male | 54 | Methotrexate 20 mg once weekly, Folic Acid | 4.22 moderate disease activity |
| 1003 | Seronegative rheumatoid arthritis | Female | 35 | Sulfasalazine 2 g daily, Vitamin D, Paracetamol | 4.21 moderate disease acitvity |
| 1004 | Rheumatoid arthritis. | Female | 44 | Methotrexate 10 mg, Hydroxychloroquine, Rituximab (previous cycle 5 months ago) | 3.09 (low disease activity) |
| 1005 | Seropositive rheumatoid arthritis (both rheumatoid factor and anti-CCP positive) | Male | 60 | Sulfasalazine Hydroxychloroquine, Methotrexate, Rituximab | 4.21 moderate disease acitvity |
| 1006 | Seropositive rheumatoid arthritis. | Female | 61 | Methotrexate, Rituximab | 3.9 (moderate disease activity) (7/3/17) |
| 1007 | Rheumatoid arthritis, Diagnosed 1998, Rheumatoid Factor/Anti-CCP+ve | Female | 52 | Etanercept since 18/07/2016, Methotrexate 22.5mg once weekly form this week, Folic Acid 5 mg six days a week, Omeprazole 20mg prn, Paracetamol, Fluoxetine 40mg od | 1.62 (remission) |
| 1008 | Psoriasis Arthritis (monoarthritis with psoriasis) | male | 41 | None | No active disease |
| 1009 | Rheumatoid arthritis, Anti-CCP positive. | Female | 36 | Methotrexate, Paracetamol Ibuprofen. Will re start Hydroxychloroquine, Folic Acid and Codeine | 4.54 (moderate disease activity) |
| 1010 | Rheumatoid factor borderline, anti-CCP positive rheumatoid arthritis. | Female | 47 | Iron tablets, stopped MTX 13 months ago, Vitamins | 1.63 (low disease activity) september 2016 score |
| 1011 | Seronegative erosive rheumatoid arthritis. | male | 75 | Tocilizumab, Methotrexate, Sulfasalazine, Hydroxychloroquine, | 2.53 (Remission, low disease activity) |
| 1012 | Rheumatoid arthritis. | Female | 38 | Methotrexate 10 mg once weekly, Folic acid, Sertraline, Rituximab | not documented |
| 1013 | Seropositive rheumatoid arthritis. | male | 68 | Tocilizumab, Methotrexate, Sulfasalazine, Hydroxychloroquine | 4.96 (moderate disease activity) |
| 1014 | Seronegative rheumatoid arthritis | Female | 60 | Tocilizumab, Hydroxychloroquine, Atenolol,Ramipril,Atorvastatin,Warfarin, Amitryptiline, Amlodipine, Aspirin | 3.25 (moderate severity) |
| 1015 | Rheumatoid factor positive, anti-CCP positive rheumatoid arthritis. | Male | 65 | Rituximab (as demonstrated three months later, not succesful) (previous Methotrexate, Leflunomide) | 4.2 (moderate disease activity) |
| 1016 | Rheumatoid arthritis | Female | 74 | Infliximab, Methotrexate | low disease activity |
| 1017 | Rheumatoid arthritis | Female | 57 | Rituximab Methotrexate | 1.65(low disease activity |
| 1018 | Rheumatoid arthritis | FEMALE | 62 | Leflunomide, Tocilizumab | Severe RA but controlled |
| 1019 | Seropositive rheumatoid arthritis with mild interstitial lung disease | male | 76 | Infliximab, Methotrexate | 5.6 high disease activity ( DAS score 5.6, 26th july 2017) |
| 1020 | Seropositive rheumatoid arthritis | male | 62 | Tocilizumab Methotrexate | 2.03 low disease activity |
| 1021 | Seropositive, anti-CCP positive rheumatoid arthritis | female | 60 | Leflunomide, Tocilizumab | 6.5(high disease activity) |
| 1022 | Psoriatic arthritis. | Female | 28 | Methotrexate, Folic acid | 4 tender joints, none swollen |
| 1023 | Undifferentiated seronegative inflammatory arthritis | male | 49 | Sulfasalazine | 2 tender joints. None swollen |
| 1024 | Seropositive erosive rheumatoid arthritis with a palindromic onset. | Female | 54 | Methotrexate, Leflunomide, Hydroxychloroquine for decision for Rituximab | 3.73 (moderate disease activity) |
| 1025 | Seronegative rheumatoid arthritis | Female | 56 | To start Methotrexate, Folic Acid | 3.61 (calculated on 3 variables moderate disease activity) |

**Suppl Table 1**
